# The neurovascular coupling in the attention during visual working memory

**DOI:** 10.1101/2023.09.28.559891

**Authors:** Hao Zhang, Yiqing Hu, Yang Li, Dongwei Li, Hanli Liu, Xiaoli Li, Yan Song, Chenguang Zhao

**Affiliations:** School of Systems Science, Beijing Normal University, Beijing, 100875, China; Center for Cognition and Neuroergonomics, State Key Laboratory of Cognitive Neuroscience and Learning, Beijing Normal University at Zhuhai, 519087 China; Chinese Institute for Brain Research, Beijing 102206, China; State Key Laboratory of Cognitive Neuroscience and Learning, Beijing Normal University, Beijing, China; Department of Applied Psychology, School of Arts and Sciences, Beijing Normal University, Zhuhai, China; Department of Bioengineering, the University of Texas at Arlington, Arlington, Texas, USA; International Academic Center of Complex Systems, Advanced Institute of Natural Sciences, Beijing Normal University, Zhuhai, China

**Keywords:** attention, EEG-fNIRS, visual working memory, neurovascular coupling

## Abstract

How to focus attention during visual working memory (vWM) depends on one’s ability to filter out distractors and expand the scope of targets. Although the spatiotemporal properties of attention processes in WM are well documented, it is still unclear how the mechanisms of neurovascular coupling (NVC) between electroencephalographic (EEG) signals and hemodynamic activity of attention during vWM. To investigate the NVC mechanism underlying attention during vWM, we recorded simultaneous functional near-infrared spectroscopy (fNIRS) and EEG data when humans were performing cued change-detection tasks. The multimodal data showed that the control and scope processes during vWM were involved in similar temporal profiles of frontal theta event-related synchronization (ERS) and posterior contralateral delay activities (CDA), and revealed similar distributions of hemodynamic activation within the frontal eye fields (FEF) and superior parietal lobule (SPL). These task-related features have a common NVC outcome across individuals: the higher EEG features (theta ERS or CDA amplitude), the greater the increment of local oxygenated hemoglobin (HbO) signals within the FEF and SPL. Moreover, when distractors should be filtered out, EEG-informed NVC is involved in a broader range of brain regions in the frontoparietal network (FPN). These results provided unique neurovascular evidence for the mechanisms of attention scope and control in vWM. Interestingly, there might be a negative relationship between behavioral metrics and theta-informed NVC strengths within the FEF for attention control. On a dynamic basis, the NVC features had higher discriminatory power for predicting behavior than EEG features and fNIRS features alone. Together, these results highlight what multimodal approaches can advance our understanding of the role of attention in vWM and how the fluctuations of NVC are associated with actual behavior.

## Introduction

In daily life, continuous visual information is maintained and manipulated over short intervals in visual working memory (vWM) to guide ongoing behavior [1]. We encode, maintain, and retrieve with our vWM largely representing the involvement of attention [2]. A limited number of available vWM “slots” [3, 4] or limited resource allocation [5, 6] requires attention processes that govern access to the mnemonic processes [7, 8]. Various processes of attention dynamically interacting with vWM have been studied, including vigilance [9], selective attention [10], and sustained attention [11].

Prior work has shown that individuals with higher vWM capacity demonstrate higher attention performance [12]. At the within-subjects level, attention and vWM covary on a moment-to-moment basis and lapse together [13]. Extensive theoretical work on information processing has long been developed on the role of attention in vWM [14, 15]. According to independent contributions to vWM performance, attention processes can be subcategorized into scope and control, which gradually become the different basic conceptualizations of attention in vWM [16, 17]. The scope of attention is an important aspect that is theoretically and empirically related to the storage of vWM and its changes affect the number of items that can be maintained in vWM [14, 18]. Also, an individual’s ability to efficiently filter relevant from irrelevant information contributes to vWM capacity [14, 19, 20]. Attention control is essential to filter out surrounding distractors when individuals need to prioritize to-be-remembered information [21, 22].

Behavioral studies have already demonstrated that the scope and control measures of attention reveal a significant positive correlation [16, 17, 23]. The neural explanation is that two attention processes in vWM have one putative source of the frontoparietal network (FPN), it is related to bias and maintenance of the goal-relevant information within certain scopes [24, 25]. The scope and control of attention might have interrelated spatial-temporal characteristics that have been well documented by electroencephalographic (EEG), functional near-infrared spectroscopy (fNIRS), or functional magnetic resonance imaging (fMRI). More specifically, the strength of the hemodynamic response [26–28] and the amplitude of EEG’s contralateral delay activity (CDA) [29–31] in the posterior parietal cortex (PPC) can be used to track the number of items maintained in vWM. Several EEG studies have shown that theta oscillations over the prefrontal cortex (PFC) are typically associated with the scope of vWM, as well as attention control [32, 33]. The prefrontal areas showed increased blood oxygen level-dependent response to increases in vWM scope or attention control [34, 35]. However, the majority of previous work has used unimodal recording or parallel analysis methods, making it difficult to provide a more comprehensive insight into the function of attention during vWM. For example, the hemodynamic activities only indirectly reflect the underlying patterns of electrical neural activity, mainly because of slow neurovascular interactions [36, 37]., which are known as neurovascular coupling (NVC). Different brain regions or different populations have different NVC relationships [38], which has implications for understanding disrupted NVC that occurs during neuropsychiatric disorders [39] and aging [40]. The NVC has been related to cognitive function including perception [41, 42], motor control [43, 44], and attention [45–47]. Especially, the NVC of anticipatory attention has an important influence on behavioral performance due to its substantial flexibility in its polarity and strength variability [47]. However, the NVC mechanism underlying the attention during vWM is still unclear.

In the current study, we therefore aimed to first identify the NVC mechanism underlying the attention control and scope during vWM then explore the associations between fluctuations of NVC and actual behavior. We recorded simultaneous fNIRS and EEG data, which allowed for assessment of NVC in a flexible, no-interference, and low-cost way. A vWM paradigm with three conditions [31] was used to isolate attention activity in vWM that occurs between presentations of the to-be-remembered array and a subsequent probe display. We tracked the changes in biological and behavioral metrics by operating on the number of targets or distractors to isolate distinct control and scope activities. We chose the prefrontal theta and parietal CDA—two robust vWM biomarkers, as main temporal features [48]. The spatial resolution of fNIRS was leveraged to functionally localize subject-specific regions of interest (ROIs) in the PFC and bilateral PPC. These experimental and analysis frameworks have the potential to aid our understanding of the NVC mechanism of attention during vWM.

## Results

The methods and material of this study are available at the Open Science Framework (http provided after publication)

### Behavioral results

To measure the control and scope of attention in vWM, we used a change-detection paradigm [31], which required the participants to remember the orientations of red bars in the cued hemifield (Fig. 1A). After a 900 ms delay, participants indicated whether the probe was the same as the orientations initially presented in the cued hemifield. For descriptive purposes, between-condition comparisons relative to baseline condition are named “attention scope” and “attention control” processes respectively. More specifically, when compared with the “2 targets” condition (abbreviated as 2T), increasing the set size of the distractor (blue bar) in the “2 targets + 2 distractors” condition (abbreviated as 2T2D) can be attributed to differences in control of attention and not to task difficulty per se, and increasing the set size of the target in the “4 targets” condition (abbreviated as 4T) was used to measure attention scope. We calculated the vWM performance using a standard formula[49]. The K value denoted the number of targets maintained in vWM from serval to-be-remembered N items. The results showed a ceiling effect (K = 1.942 ± 0.025 VS. N = 2) for the 2T condition. The indices for attention scope were chosen as the maximum K value (2.117 ± 0.488) through comparison between the 4T and 2T conditions. A higher scope value denoted a larger attention scope. The indices for attention control (1.749 ± 0.207) were defined as the difference in K values between the 2T2D condition and 2T condition[23, 49]. A higher attention control value indicated the better ability to filter out distractors. Our results showed that the indices for scope were significantly correlated with the indices for control (Pearson’s r = 0.406, p = 0.001, Fig. 1B), highlighting that individuals with larger scope have better attention control to filter out distractors. This result is consistent with previous research [22].

**Fig. 1.**
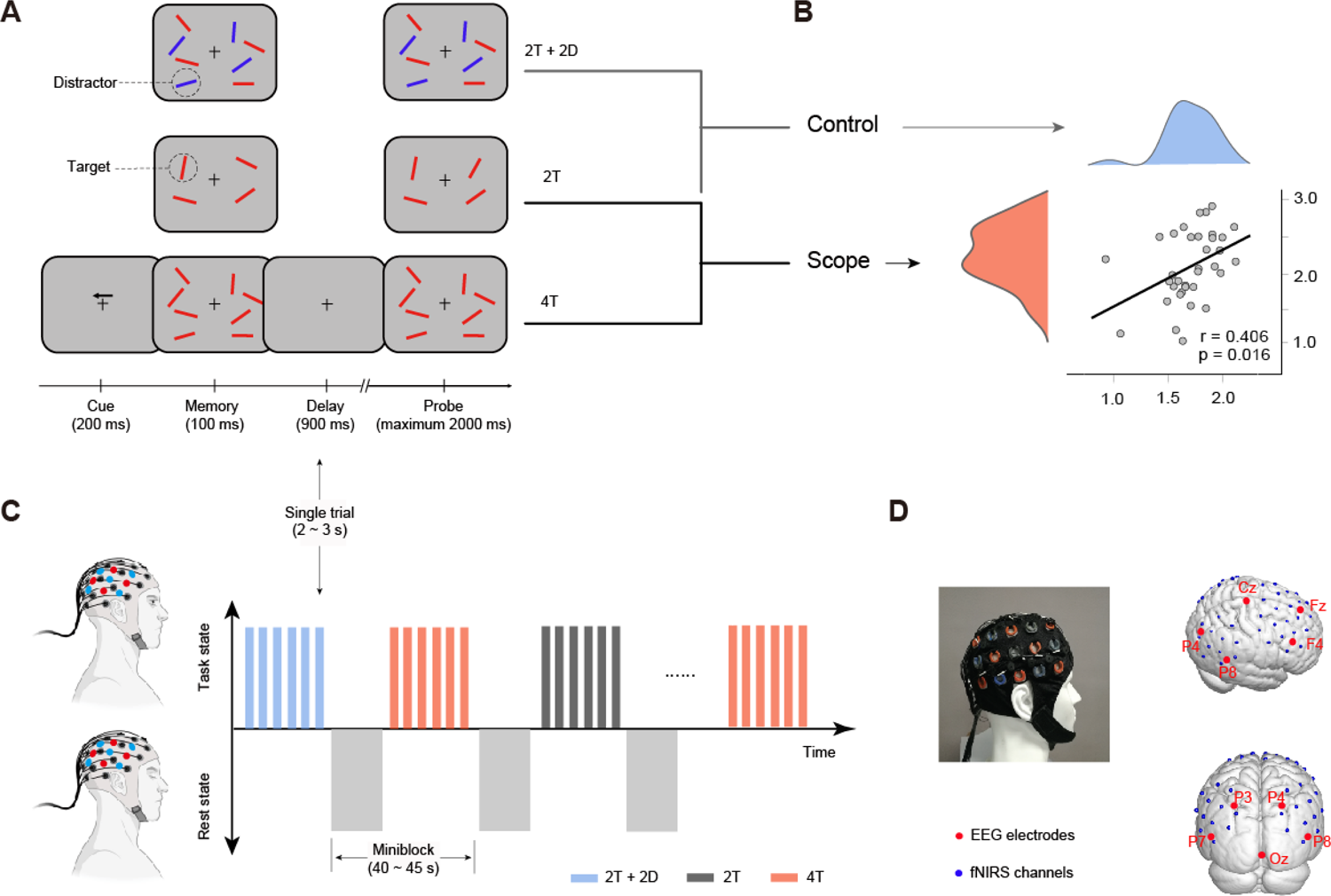
Experimental design and simultaneous data acquisition. (**A**) vWM task: two targets (2T), two targets and two distractors (2T+2D), and four targets (4T), three conditions were designed to isolate the control and scope of attention during vWM by comparing the baseline conditions (2T). (**B**) Correlation of control and scope: a robust positive relationship between attention control (blue, bottom axis) and attention scope (red, right axis). (**C**) Miniblock design nesting rest state: by introducing a closed-eye period between miniblocks, a transition between task and rest states is achieved while preventing temporal overlap of prior hemodynamic responses. (**D**) Sensor localization: a photograph of the sensor cap and side/rear views of EEG electrodes (red) and fNIRS channels (blue), covering regions of interest, such as the prefrontal cortex and posterior parietal cortex.

### Spatiotemporal features in the PFC

Simultaneous EEG and hemodynamic signals were collected and analyzed using a miniblock-design paradigm (Fig. 1C). As shown in Fig. 1D, we measured the concentration changes in oxygenated hemoglobin (ΔHbO) and deoxygenated hemoglobin (ΔHbR) in the PFC and PPC, as well as whole-brain EEG data. It is reported that the frontal theta is a prominent EEG feature in the attention or WM task, which is theorized to play important roles in either the control or scope of attention representations [16, 17]. We estimated this modulation of frontal theta power during the retention of vWM by calculating the theta event-related synchronization (ERS). Consistent with many previous studies [32, 50, 51], the amplitudes of the theta ERS in the three conditions were increased by approximately 100–400 ms after memory onset (Fig. 2A; FDR p < 0.050, 2-tailed), suggesting that theta power served as a possible carrier frequency for vWM processes [52]. To further segregate the control-related and scope-related theta activities, we contrasted baseline conditions (2T) with two others. As shown in Fig. 2A, the theta ERS significantly increased in the 2T2D condition (from 120 to 390 ms) and in the 4T condition (from 140 to 400 ms) relative to the 2T condition (cluster-based permutation test: p < 0.050, two-tailed). These differences shown in topographic maps were averaged over significant time points. The scalp distributions of increasing theta ERS in both 2T2D and 4T conditions were mostly expressed over the same prefrontal electrodes (F3, F_Z_, and F4). No significant difference in theta ERS between 2T2D and 4T was found (p > 0.371; BF_10_ = 0.101). Our EEG results showed similar scalp distributions and time course profiles for control and scope of attention in vWM.

**Fig. 2.**
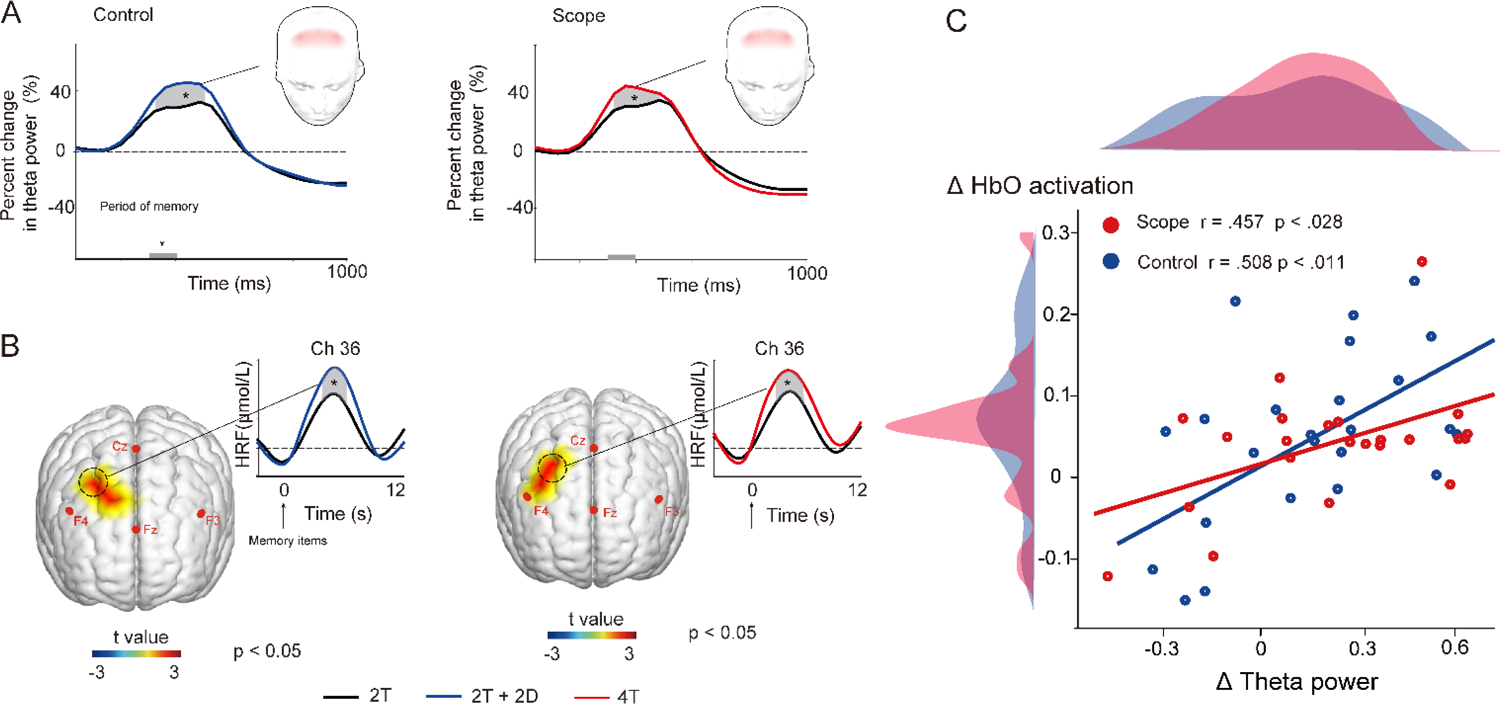
Spatiotemporal features in the PFC. (**A**) frontal theta ERS: a consistent increase at approximately 120 to 400 ms post memory onset, and the red-shaded region indicates significant electrodes (frontal view). The gray-shaded regions, marked with asterisks, indicated statistically significant time intervals. (**B**) hemodynamic activation: The brain map highlights changes in ΔHbO responses within brain regions exhibiting statistically significant t values (frontal view) compared to the baseline condition. The ΔHbO signal response exhibits a similar temporal pattern. (**C**) NVC analysis across subjects: The correlation between relative changes in Δtheta power and ΔHbO within ROIs demonstrated a positive correlation, with the shaded areas on the left and top axes indicating the distribution of values for each feature. * p < 0.05.

We then focused on the difference in hemodynamic activity (as indicated by ΔHbO levels) related to the scope and control of attention. For the attention control, paired t-tests showed that significant increases in hemodynamic response were found in the PFC near the F4 electrode and defined control ROI (channels 36 and 32) in Fig. 2B; FDR p < 0.050, 2-tailed). We also found a significant increase in the ΔHbO response for the attention scope in the partially overlapping regions of the control and defined scope ROI in Fig. 2B (channels 36 and 31; FDR p < 0.050, 2-tailed). Structural MR images with extra anatomical landmarks (vitamin E capsules) were used to project the fNIRS channels onto the cortical surface through the NIRS_SPM package [53] and determine the anatomical localization of each fNIRS measurement channel. The results showed that the anatomical localization of scope ROI and control ROI were both in the frontal eye fields (FEF) (Table S1). The overlapping channels (channel 36) were used to show the time course of memory response by general linear model (GLM) analysis [54]. The attention scope led to a significant increase of ΔHbO from 4.1 s to 7.6 s locked to memory display (cluster-based permutation test: p < 0.050, Fig. 2B). Similar to attention scope, a significant increase from 3.8 s to 7.7 s was observed for the attention control. No significant difference of ΔHbO between 2T2D and 4T was found (ps > 0.241; BF_10_ < 0.207).

We performed between-subject correlation analyses to quantify the NVC between theta ERS and ΔHbO activation. As shown in Fig. 2C, the difference in theta ERS between the corresponding condition and baseline was positively correlated with the hemodynamic difference within functional ROIs (for attention control: r = 0.508 and p < 0.011; for attention scope: r = 0.457 and p = 0.028, 2-tailed). As mentioned above (Fig. 2), both theta ERS and ΔHbO activations have similar spatiotemporal characteristics in the PFC and NVC between the individual levels, which suggests that scope and control processes might share common functional mechanisms in the PFC.

### Spatiotemporal features in the PPC

We used CDA as an EEG marker of vWM in humans to track the change in active storage during the retention of vWM [30]. Previous studies have demonstrated that the contralateral effect (contralateral to target > ipsilateral to target) was observed over the PPC in fMRI and fNIRS studies [55–57]. Consistent with many previous studies [29–31], our results showed that a sustained, negative potential with a posterior scalp distribution persists during the delay period (Fig. 3A; FDR p < 0.050, 2-tailed). EEG CDA amplitudes were compared across conditions to assess the scope and control of attention. As shown in Fig. 3A, the amplitude of EEG CDA rose from 425 to 900 ms as attention scope increased (Fig. 3A; cluster-based permutation test: p < 0.050, 2-tailed), which is consistent with the fact that the posterior CDA amplitude was sensitive to mnemonic load [58]. For attention control, our results showed that the CDA amplitude also significantly increased from 477 to 900 ms (cluster-based permutation test: p < 0.050, two-tailed). The topographic maps of these differences over significant time points have shown similar scalp distributions, where they were mostly expressed over the same posterior parietal electrodes P3 (paired P4) and P7 (paired P8). No significant difference of EEG CDA between 2T2D and 4T was found (p = 0.114; BF_10_ = 0.217). Our EEG results showed that control and scope of attention are involved in CDA changes with similar scalp distributions and time course profiles.

**Fig. 3.**
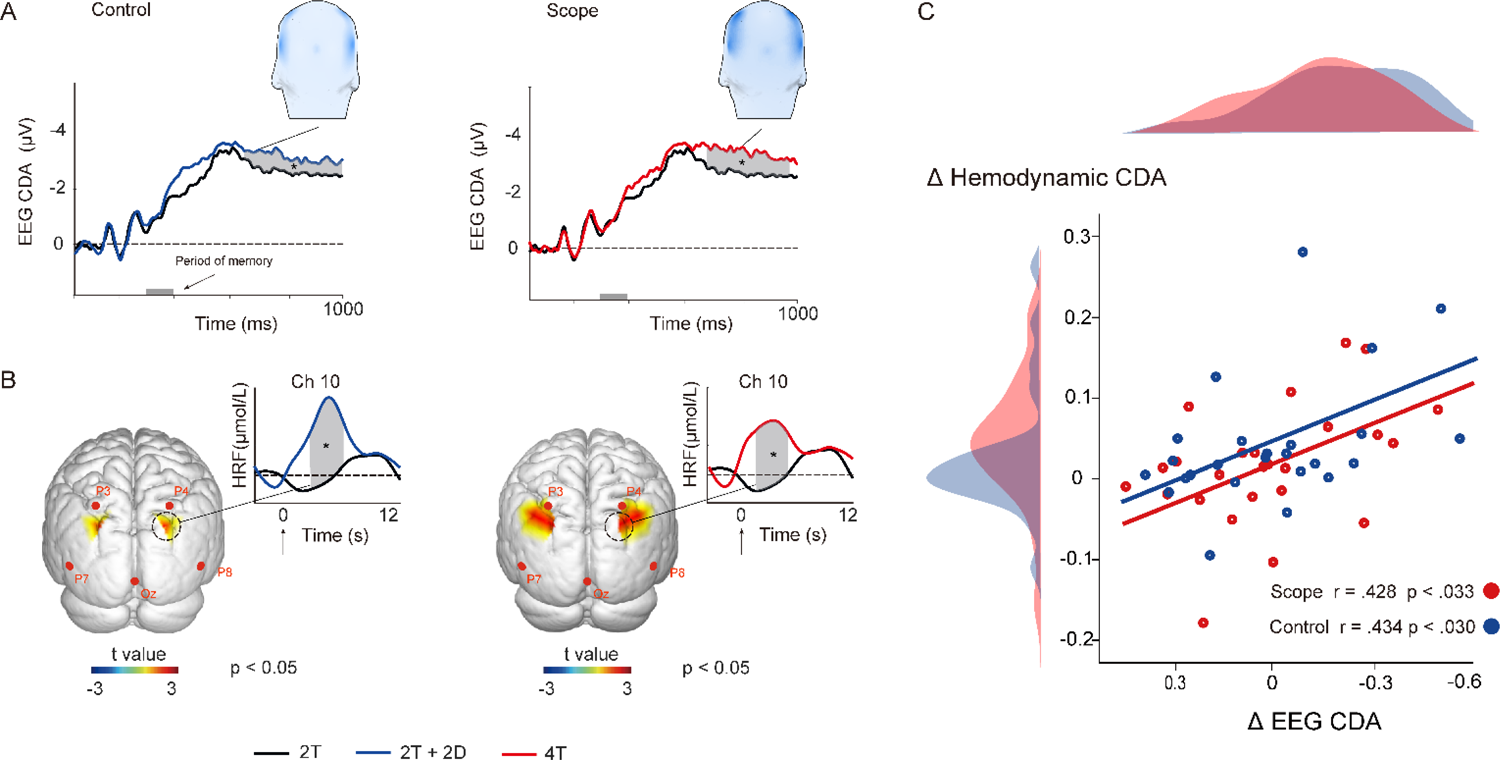
Spatiotemporal features in the PPC. (**A**) EEG CDA: variation during the delay period revealed sustained negative potential changes (amplitude axis reversed), with the blue-shaded region indicating significant electrodes (posterior view). The gray-shaded regions, marked with asterisks, indicated statistically significant time intervals. (**B**) hemodynamic CDA: The brain map highlights changes in contralateral effects within brain regions exhibiting statistically significant t values (posterior view) compared to the baseline condition. The ΔHbO signal response exhibits a similar temporal pattern. (**C**) NVC analysis across subjects: The extracted characteristics from ΔEEG CDA and Δ hemodynamic CDA demonstrate a positive correlation for attention control and scope. with the shaded areas on the left and top axes indicating the distribution of values for each feature. * p < 0.05.

To investigate metabolic features related to attention in vWM, we quantified the hemodynamic CDA by using a similar approach to the EEG analysis for each pair of symmetrical PPC channels. For the attention control, paired t-tests showed that significant hemodynamic CDA increases were found in the PPC near the region between the P3 and P7 electrodes (channels 10 and 9 in Fig. 2B; FDR p < 0.050, 2-tailed). We also found a significant hemodynamic CDA increase for the attention control in the overlapping regions (channels 10 and 20) and extra regions (channels 9 and 21), as shown in Fig. 3B (FDR p < 0.050, 2-tailed). The results (Table S2) showed that the anatomical localization of the bilateral channels 10 (paired 20) and 9 (paired 21) were projected into the bilateral superior parietal lobe (SPL). The waveform of hemodynamic CDA in the overlapping channels (channel 10) showed that memory item led to a significant increase of hemodynamic CDA from 2.4 s to 7.1 s for the attention scope (cluster-based permutation test: p < 0.050, Fig. 3B). Similarly, a significant increase from 2.0 s to 6.6 s was observed for the attention control. No significant difference of ΔHbO between 2T2D and 4T was found (ps > 0.211; BF_10_ < 0.307).

As illustrated in Fig. 3C, the difference in hemodynamic CDA activation was closely and positively correlated with the difference in EEG CDA amplitude across all participants (for attention control: r = 0.446 and p < 0.040; for attention scope: r = 0.563 and p < 0.020), which suggested that the more increase of posterior CDA amplitudes was, the more increase in local hemodynamic changes observed nearby at the corresponding electrodes. Note that the EEG CDA amplitudes were negative, and the y-axis in Fig. 3A is plotted negative-up. To render its polarity consistent relative to the trend of CDA, we show the negative-right axis of ΔCDA in the correlation scatter plot. In addition, we analyzed parietal contralateral alpha power suppression (8-12 Hz), which was also found during vWM tasks. No significant difference was found across the three conditions (ps > 0.577). We also analyzed the ΔHbR concentration by using the same methods as those used for the ΔHbO data. Neither a significant activation nor a reliable correlation between the ΔHbR concentration and corresponding EEG features was found for any of the conditions.

### EEG-informed NVC

Thus far, we have outlined the basic NVC for scope and control of attention at between-subject levels: EEG activities (CDA or theta power) are seemly accompanied by the resulting metabolic variation. This qualitative NVC analyses under simplifying assumptions that the NVC is constant for every individual throughout the experiment. However, attention is not constant and varies with time[59]. We further explored the NVC mechanisms underlying the control and scope of attention by utilizing EEG-informed NVC analysis, which can move beyond the above correlational analyses by monitoring and assessing the trial-by-trial variance in inter-subject activity. We hypothesized that posterior CDA and frontal theta ERS have links with the ΔHbO concentration variation at within-subject levels, resulting in observed significant EEG-informed NVC. To quantify the NVC, EEG-informed NVC analysis modeled the association of hemodynamic activity and WM-derived EEG characteristics by using GLM, and t-contrasts showed the significance of the coefficients related to the WM-derived EEG regressors in the GLM at the within-subject level. For the 2T2D condition, the frontal theta ERS was positively correlated with ΔHbO in the right FEF, right SPL (Fig. 4A, first row, FDR p < 0.05). That is, front theta ERS was more elevated when these brain regions were more active on a trial-by-trial basis. The bilateral SPL and the FEF showed significant NVC with posterior CDA amplitude (Fig. 4A, second row, FDR p < 0.05, 2-tailed). For the 4T condition, regions showing significant NVC of frontal theta ERS included only FEF and not SPL (Fig. 4B, first row, FDR p < 0.05, 2-tailed). The coordinates of the regions showing significant NVC are listed in Table S3. For EEG CDA from the PPC, areas showing positive NVC included regions within the right SPL (Fig. 4B, second row, FDR p < 0.05, 2-tailed). That is, on a trial-by-trial basis, these brain regions showed increased ΔHbO activity when the CDA amplitude increased. As expected, increased use of posterior CDA or frontal theta power were both accompanied by the increased adjacent ΔHbO for the 2T2D and 4T conditions. Interestingly, our results showed that NVC under the 2T2D condition is involved in a broad range of brain regions, some of which are part of the FPN. In contrast, no significant NVC across regions was observed in the 4T condition (ps > 0.358). This difference in activated regions of NVC provided the dissociation of the mechanisms of scope and control of attention during vWM, suggesting that NVC can be characterized by substantial flexibility in its polarity and spatiotemporal variability [60–62].

**Fig. 4.**
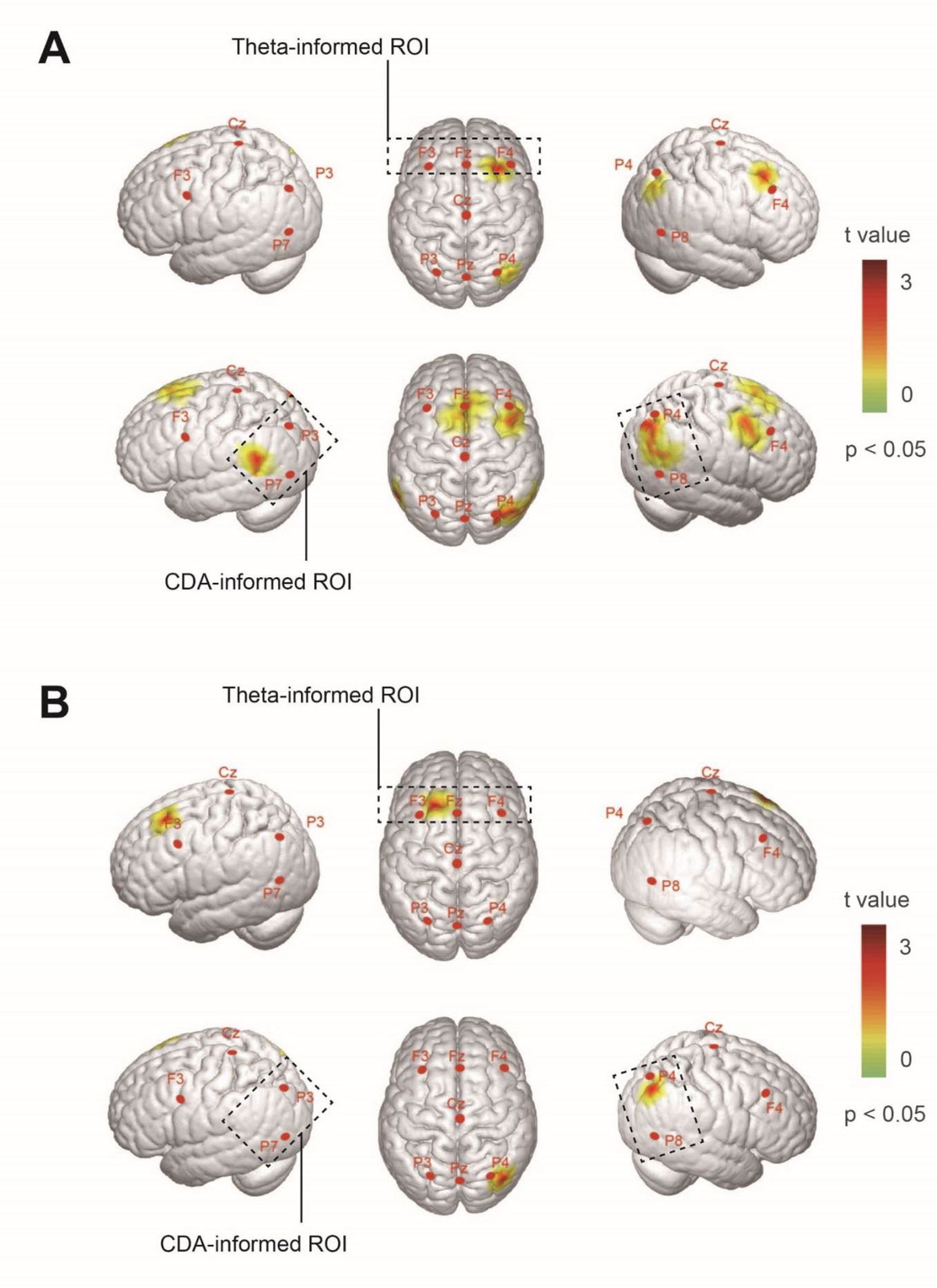
The EEG-informed NVC. (**A**) EEG-informed NVC for attention control. the t map shows the significant theta-informed NVC (upper panel) in the middle frontal gyrus and SPL and CDA-informed NVC (bottom panel) in the bilateral SPL and middle frontal gyrus. (**B**) EEG-informed NVC for attention scope. t map shows the significant theta-informed NVC (upper panel) in the middle frontal gyrus, and CDA-informed NVC (bottom panel) in the bilateral SPL.

### Dynamic NVC and K value

We further proposed a dynamic EEG-informed algorithm to estimate NVC over the integration of temporally sensitive EEG information and spatially sensitive fNIRS information. First, we transferred the analytical framework to a novel dynamic behavioral performance and NVC built on miniblock experimental designs. The 10 s closed-eye resting state was interleaved between miniblocks to speed up the return of the hemodynamic signal to prevent a temporal overlap of the prior hemodynamic response. A simple illustration in Fig. 5A shows that the ΔHbO concentration was reduced to the prior baseline after recovery of the closed-eye resting state (see SI for details).

**Fig. 5.**
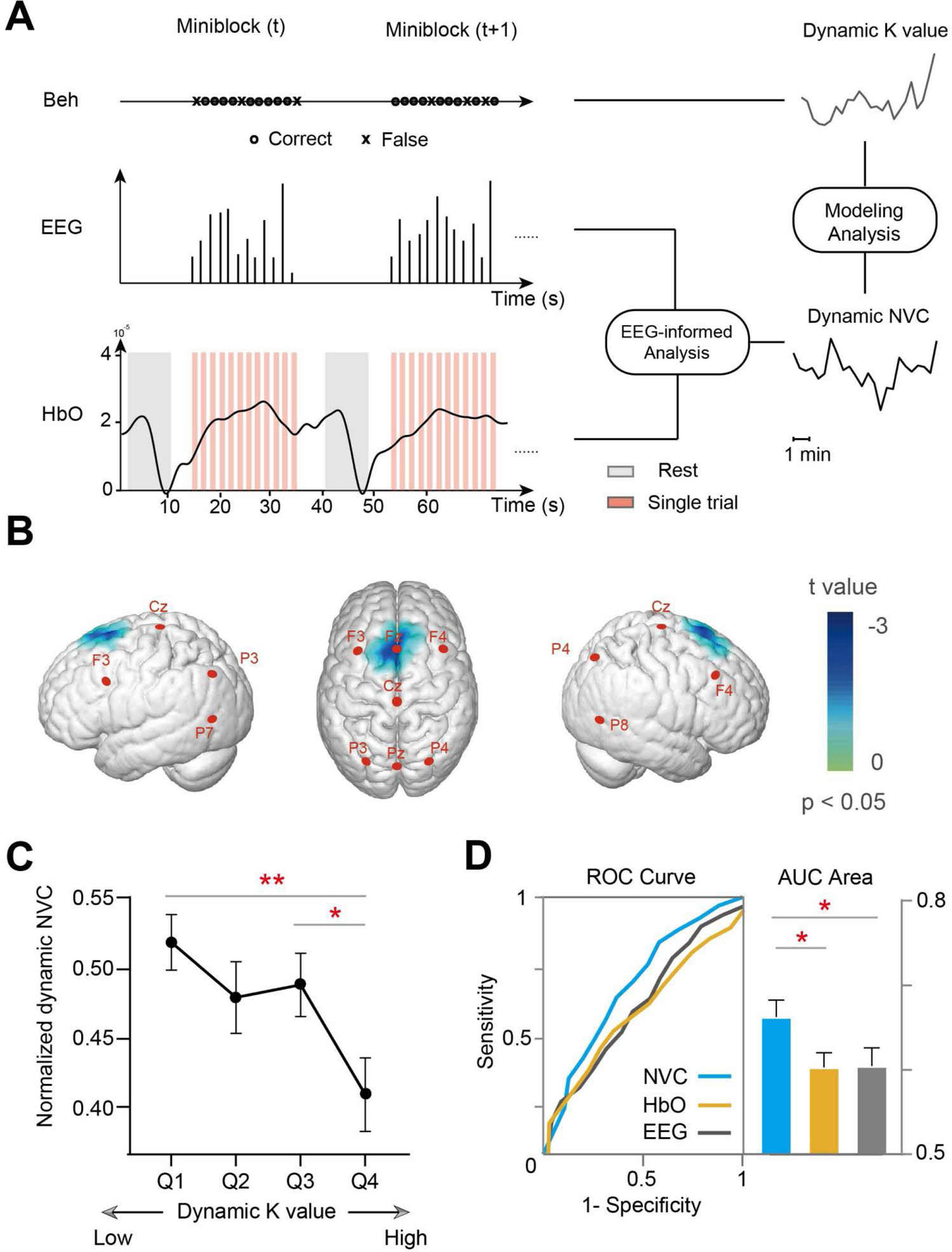
The relationship between NVC and behavior. (**A**) The upper row shows behavioral changes in each miniblocks, the middle row shows quantified EEG features, and the bottom row presents the hemodynamic response alternating between task and rest. Then, EEG-informed analysis investigates the dynamic NVC. The dynamic K values are integrated with dynamic NVC, leading to the modeling analysis. (**B**) The coefficient brain map revealed a significant cluster in the FEF, indicating a negative correlation between theta-informed NVC and dynamic behavior metrics. (**C**) The dynamic K values were divided into four quadrants, there was a gradual reduction in the normalized dynamic NVC with increasing dynamic K values. (**D**) left panel: ROC analysis predicting dynamic K values from the theta-informed NVC (blue), standalone ΔHbO (yellow), and standalone EEG (gray). right panel: area under the curve (mean ± standard error) of the ROC analysis. * p < 0.05; ** p < 0.01

Moreover, we conducted a modeling analysis of how the dynamic NVC can influence dynamic behavior as measured by the K value. The GLM included EEG-informed NVC (theta-informed or CDA-informed) features as the regressor and used the dynamic K value in different conditions as the data to be regressed for each individual (see the pipeline in Fig. 5A). To visualize our results, we plotted a coefficient brain map with a statistical t value at a typically thresholded FDR value p < 0.05 (see Fig. 5B). For frontal theta-informed NVC, the FEF showed a significant cluster in the 2T2D condition. No significant cluster was found in other conditions or other EEG-informed NVC (p > 0.217; BF_10_ < 0.301). The negative coefficient indicated that the frontal theta ERS was more closely coupled to adjacent ΔHbO signals, and the smaller dynamic K value was.

We also performed a correlation analysis between the dynamic NVC with the dynamic K value. It showed that the K values were smaller in miniblocks with a weaker NVC of theta ERS over miniblocks across subjects(Spearman’s r = −0.113, p = 0.014, 2-tailed). We further calculated the average every miniblock NVC for each quartile to confirm the relationship between frontal theta NVC and the K value. Repeated-measures ANOVA of NVC showed that the normalized NVC decreased with an increase in the K value in the 2T2D condition (F_3, 75_ = 4.450, p = 0.007). The results in Fig. 5C show that the NVC in the fourth quartile was significantly smaller than that in the first quartile (t = 3.599, FDR p = 0.002, Cohen’s d = 0.750) and third quartile (t = 2.851, FDR p = 0.009, Cohen’s d = 0.594). Similarly, no significant difference was found among the quartiles (ps > 0.239) in the other condition.

The receiver operating characteristic (ROC) analysis was used to compare the sensitivity and specificity of various WM-related metrics to predict the dynamic performance for the 2T2D condition. We evaluated the performance of the GLM analysis concerning (EEG, ΔHbO, and NVC metrics), and showed results in Fig. 5D. The one-way repeated ANOVA analysis indicated a significant effect of the model type (F_2,44_ = 3.339, p = 0.045) on the area under the curve AUC values that represented the sensitivity and specificity of the model. The posthoc test indicated that the NVC-based model achieved the highest AUC that was significantly higher than the EEG-based (t = 1.929, p = 0.033, Cohen’s d = 0.402) and ΔHbO (t = 2.330, p = 0.015, Cohen’s d = 0.486) model. Finally, when fitting the model predicting the performance of attention scope, no significant difference in the AUC was found among three WM-related metrics (ps > 0.271). These results indicated that the NVC for the control of attention can be predictive of behavioral outcome and therefore is likely to play an important role in cognitive processes related to memory and attention. We also analyzed the data based on reaction times and the variance in reaction times using the same methods, but no significant results were found in any of the conditions (ps > 0.427).

## Discussion

In this study, we investigated the spatiotemporal features and the corresponding NVC for attention scope and control during vWM through both temporally sensitive EEG markers and spatially sensitive hemodynamic activation. The main findings were as follows: (1) The multimodal results showed that the control and scope processes in vWM evoked similar temporal profiles of frontal theta ERS or posterior CDA and similar distributions of hemodynamic activation within the FEF and SPL. Crucially, we provided converging evidence that the control and scope processes share a similar NVC mechanism at both the trial and trait levels, except that the attention control involved a broad range of NVC within FPN. (2) The fluctuations of theta-informed NVC within SPL can be predictive of the performance to filter out distractors. There might be a negative relationship between behavioral metrics and dynamic NVC strengths. Moreover, theta-informed NVC might indeed be potential candidates for predicting behavioral performance involving attention control. In sum, these results highlight that multimodal approaches can advance our understanding of the NVC mechanism of attention in vWM.

### The NVC with frontal theta activities

Frontal theta oscillations play a causal role in facilitating increases in vWM capacity, which are thought to have links with frontal theta power [32, 63] or with theta synchronizations [64]. Our oscillatory results showed that there was a greater increase in frontal theta power as target items or distractor items increased. This is consistent with the idea that increased frontal theta power during the period of vWM representation played a positive role in the increasing demand for attention [50, 65]. The synchronous fNIRS results revealed that the increased activations in the right FEF were observed for attention control and scope. It might mirror the attempt to modulate attention allocation to increased target items or distractor inhibition [66]. Interestingly, we found that a positive relationship between increased Δtheta ERS and increased ΔHbO activation appeared for attention control and attention scope processes. This NVC may be explained by the fact that FEF is the likely source location for vWM-related frontal theta activity [67]. More specifically, individuals have more increased frontal theta activities and adjacent cortical hemodynamic responses when demands of attention scope or control are increased in vWM tasks. It is consistent with previous ideas that the NVC between hemodynamic responses and theta oscillations plays a critical role in performing vWM tasks [68].

On a trial-by-trial basis, when distractors should be filtered out, the frontal theta ERS was more elevated when these brain regions in the right FEF and right SPL were more active. In contrast, when there were no distractors but rather extra targets, the NVC of frontal theta ERS included only the FEF. A possible explanation for this might be that the control of attention might be associated with the widespread NVC. As far as we know, theta-band oscillations are proposed to serve as the carrier frequency for attention control [69, 70]. In this case, we suggested that the hemodynamic activities of these two regions could be coupled with frontal theta activities.

### The NVC with posterior CDA

The CDA amplitudes within PPC increased during the retention period (approximately 400 to 1000 ms) when the set size of distractors or the scope of targets increased. This finding was consistent with previous studies using the changes in CDA amplitude to track and compare the number of items maintained in vWM [30]. From synchronously observed fNIRS results (Fig. 3), hemodynamic CDA originates from the contralateral effect in the PPC to the visual field of the target, which was reported in previous fNIRS/fMRI studies using the lateralized ERP approach [45, 47]. The rise in hemodynamic CDA in the bilateral SPL as the scope of the target in vWM increased from two to four items, which is in agreement with the findings of previous studies [27, 71, 72]. In this study, we developed synchronous measurements and analyses for the multimodal contralateral delay activities, and for the first time calculated the relationship between electrical and hemodynamic CDA. Our results showed that, the changes in hemodynamic CDA are positively correlated with the changes in EEG CDA amplitudes in the PPC during active maintenance. Given that the set-size effects of EEG CDA are accepted neurophysiological measures for the range of items held in vWM [73], when the scope of targets increases, the hemodynamic CDA is likely to reflect the linearly related metabolic demands supporting simultaneous EEG CDA during a certain period. In this sense, the role of increased hemodynamic CDA in attention control might stem from the increased demand for oxygen, as distractors failed to filter out was maintained in vWM.

In addition, the bilateral SPL and the MFG showed significant CDA-informed NVC for attention control, but not the MFG for attention scope. Given that posterior CDA amplitude can be used to track the number of distractors inadvertently reaching vWM, it is plausible that the EEG CDA may be linked to hemodynamic activity in the frontal and parietal regions. This represents a likely mechanism of vWM processes where the PFC is engaged in attention control to filter out distractors existing in the PPC buffer for vWM.

### The relationship between NVC and behavior

Recent studies have made considerable headway on how attention guides behavior in this dynamic way when completing a whole cognitive task [74]. The capacity to stabilize the content of attention not only varies across individuals but is also time-varying within individuals [75]. This means that the attention during the vWM task is not static, nor is it possible—or desirable—at each miniblock to simultaneously label the unique NVC calculated at the group level. Rather, failures to filter distractors are frequent, and the scope of attention fluctuates, paced by the natural speed of attention systems that govern access to vWM. The relationship between the task-related NVC strength and behavior performance was almost conducted across individuals [76], with subject-level analyses not reported. To fill this knowledge gap, we first continuously tracked the fluctuations in vWM processing at various levels of psychophysiological reactions (see Fig. 5) and then, for each miniblock, quantized the dynamic NVC between the moment-to-moment changes in brain activity, as well as continuous readouts of behavior. The results of modeling (explained behavioral data) showed that dynamic NVC of frontal theta activities within the FEF have a robust negative relationship with K values for attention control processes. Huo et al. reported the neurovascular decoupling phenomena that, in the frontal cortex of head-fixed mice, the firing rate and oscillatory power both increased as hemodynamic activities did not change during locomotion [77]. We suggest that NVC strength between frontal theta activities and ΔHbO plays an important role in controlling distractors filtered out of vWM. More specifically, the decoupling or coupling phenomenon of neural activity from hemodynamic signals might determine whether attention control is zoomed in or out in vWM. This relationship should be interpreted with caution unless the detailed NVC mechanisms linking attention control are available at the microscopic and mesoscopic levels [78]. In addition, diagnostic accuracy evaluation demonstrated that the strength of NVC had higher discriminatory power for predicting behavior than EEG features and fNIRS features alone, which is consistent with the idea of enhanced classification and decoding accuracy over a single modality in various tasks for brain-computer interfaces [79, 80].

### The NVC and attention research

The first studies [81–83] utilized simultaneous EEG-fMRI recordings during verbal WM tasks with a long-term trial (> 10 s). However, investigations dealing with attentional aspects of vWM such as scope and control processes are rare. The reason for this might be that the acoustic noise during fMRI scanning requires additional attention resources as confounding factors during an attention task [84]. The use of EEG-fNIRS has paved the way for the use of ecologically valid settings to eliminate acoustic artifacts in fMRI scanning. Some EEG-fNIRS studies have focused on verbal WM [85, 86] or vWM by parallel analysis [87, 88]. Few studies have been performed to investigate attention during vWM. The main reason is the lower certainty and more complexity of attention tasks compared with simple sensory and motor tasks [89, 90]. To exclude potential confounding effects, such as motivation, strategies, and adaptive habits, two of three conditions were combined to isolate distinct attention-related processes. Another challenge in effectively measuring the NVC mechanism is mismatching timescales between EEG and hemodynamic signals. The most popular task design among EEG-fNIRS studies is the block design, which improved the accommodation of block designs for the relatively slow hemodynamic response compared with the event-related design. However, block designs also brought a higher susceptibility to anticipation and habituation effects, which should be avoided in rapid event-related EEG studies [91]. Moreover, if the experiment duration is limited and intertrial intervals (ITIs) should persist longer than 10 s, the number of trial repetitions per condition decreases, potentially lowering the EEG signal-to-noise ratio (SNR). To address this dilemma, we first included the mixed block/event-related fMRI design allowed for a fuller characterization of time-sensitive and nonlinear activities [92]. The active eye-closed resting state could speed up the return of the hemodynamic signal to the baseline to ensure that hemodynamic variations are not artifacts caused by the prior signal. This miniblock design nested resting state has been thought to balance trade-offs between minimized crossover effects from inherently slow hemodynamic signals and pursuing a higher signal-to-noise ratio from a greater EEG response.

### Limitations and Future Work

There are still some limitations in the current study: first, although we provided converging evidence for NVC analysis at both the between- and within-subjects levels, individual difference correlations based on relatively modest sample sizes, make it challenging to build robust correlations across individuals due to possible inflated effect sizes [93, 94]. Second, a particularly significant disadvantage of EEG-fNIRS when evaluating the NVC is that it is only limited to superficial neocortical levels. Hence, the NVC in deep-seated structures such as the hippocampus or the thalamus, cannot be addressed. Last, in the change-detection design, the target and distractor lend themselves in the probe array. A model-driven approach, such as lateralized activity analysis and contrast with the baseline condition, was applied to separate attention processes from the compounds of vWM. However, the observed hemodynamic results may have been inflated by the attention evoked by the probe array rather than reflecting only attentional processes in vWM [57]. Despite these unresolved issues, our study provides novel insight into the NVC mechanism underlying the scope and control of attention. For future clinical research, the detailed NVC mechanism underlying basic cognitive processes will be an important step forward in the successful application of relevant theory and methods for measuring cognitive deficits associated with a wide variety of neuropsychiatric disorders [36, 39, 78]. Our work not only provides a more comprehensive insight into the functional activity but also a new perspective for the evaluation and prediction of behavioral performance on a dynamic basis. Individual moment-to-moment differences in filtered-out distractors can be related to the fluctuations of theta-informed NVC within FEF. This provided a promising approach for optimizing spatiotemporal resolution in the closed-loop system of cognitive brain-machine interfaces as the liberating potential of “1+1 > 2” from this multimodal technology.

## Materials and Methods

### Participants

35 participants were right-handed and had normal or corrected normal vision (21 females, age range 18.2–27.1 years, mean age 24.5 years) included in our experiment. Data from seven participants were excluded due to incomplete data or low signal quality. The experimental procedures of the study were approved by the Institutional Review Board of Beijing Normal University, and each participant signed informed consent forms.

### Working Memory Task

The visual stimuli were displayed on a 20-inch CRT monitor, with 1600 × 900 pixels and a refresh rate of 120 Hz, placed 60 cm from the participant. As shown in Fig. 1A, participants were instructed to keep their eyes fixated on the cross in the center of the screen throughout the whole experiment. A spatial arrow cue was presented at the beginning of each trial for 200 ms, instructing participants to remember the items that appeared in the left or the right hemifield. Next, memory items with a homogeneous light gray background (12 cd/m^2^, RGB: 125, 125, 125) were presented for 100 ms followed by a 900 ms delay with a blank display. Finally, the probe display was presented until a response was given or after a maximum time of 2000 ms and required participant answer quickly and accurately whether the orientation of the items that appeared in the cued hemifield had changed in the test display.

All memory items were nonoverlapping and randomly distributed within bilateral rectangular regions (4°×7.3°) centered 3° to both sides of a black central fixation point (0.5 cd/m^2^, 0.4°×0.4°). The distance between the center and the center of the bars within a hemifield was at least 2°. Each memory display consisted of two or four 2°×0.5° red bars with a random orientation between 0° and 180°. The orientations among bars within a hemifield differed by at least 20°. The items on the test display were the same as those used for the memory display in 50% of the trials. In the remaining 50% of trials, the orientation of one bar in the test display was changed at least 15° from the corresponding items in the memory display. In the baseline condition, two red bars were presented in each hemisphere. In the scope-increased condition, an increase of four red bars was presented in each hemisphere. In the control-increased condition (2T2D), two red and two blue bars were presented in each hemisphere.

### Miniblock nesting rest state design

Each experimental condition consisted of three blocks, with randomization applied across these blocks. Each block was separated by an approximate interval of 30 seconds. Within each block, there were six miniblocks, each separated by a 10 s rest period. The participants closed their eyes when the fixation disappeared. The participants were asked to open their eyes after hearing the onset beep, then a 2.2-second countdown allowed them to get ready for the upcoming stimulus (Fig. 1C). Every miniblock encompassed 12 trials, wherein participants were presented with randomly occurring memory tasks involving stimuli moving either leftward or rightward. Participants were required to discern the congruence of the presented stimuli based on cue indications and respond promptly with a keypress. Consequently, each unique experimental condition comprised a total of 3 blocks × 18 miniblocks × 12 trials = 648 trials. The entire experimental session was extended for an approximate duration of 50 minutes.

### Simultaneous data acquisition

Simultaneous EEG and fNIRS signals were recorded while the participants performed all tasks. The EEG data were recorded by using a SynAmps2 amplifier and the Curry 8.0 package (Compumedics Neuroscan, USA). fNIRS signals were recorded on an ETG-4000 system at 830 nm and 695 nm (Hitachi Medical, Japan). Thirty-two Ag-AgCl electrodes from an EEG cap were arranged according to the international 10–20 system. Four additional facial electrodes were placed at the outer canthi and 1 cm above and below the left eye to record the horizontal or vertical electrooculogram (EOG) data; two electrodes were placed at the left and right mastoid. 46-channel fNIRS measurement from one 3×5 and two 3×3 optode probe sets that covered the PFC and PPC (Fig. 1D). fNIRS signals acquired at 10 Hz and EEG signals acquired at 1000 Hz were synchronized via an external event-triggered marker generated by Psychtoolbox-3 (http://psychtoolbox.org). A linear correction was applied to the timeline of the data using the relative time of identical markers, resulting in a synchronization error of less than 0.1 s. For online EEG recording, electrode impedances were maintained below 5 kΩ for all remaining electrodes. The EEG data was digitized online and a bandpass filter of 0.01–200 Hz was applied.

### Preprocessing

Offline EEG data processing and analyses were performed using MATLAB software (The MathWorks Inc., MA, USA) with EEGLAB toolbox [95], ERPLAB toolbox [96], and custom codes. The EEG signals were 0.1–40 Hz bandpass filtered and referenced to the average of the left and right mastoids. The filtered EEG data were decomposed into independent components using the independent component analysis (ICA) algorithm. One or two components that showed the highest correlation with vertical EOG (eye blink) were removed. The EEG data were then segmented from –500 to 1200 ms concerning memory display. EEG epochs exceeding ±100 μV at any electrode or horizontal EOG exceeding ± 30 μV were automatically excluded from 0 to 500 ms around memory display onset. Data from 4 participants with a high ratio (>40%) of excluded trials were discarded from the participants’ final dataset. An average of 12.1±6.7% of trials per participant were rejected due to artifacts.

For offline NIRS data processing and analyses, light intensity changes were converted to the concentration changes in deoxy- and oxyhemoglobin (ΔHbR and ΔHbO) according to the modified Beer-Lambert law. Bad channels were detected and interpolated using adjacent channel data. Channels with the following conditions were marked as bad channels: (1) the coefficient of variation (CV) deviated from the average value by more than 90% [97]; (2) the correlation coefficient between ΔHbO and ΔHbR was not significant [98]; and (3) the power spectrum of the fNIRS signal did not show a significant peak in the heart rate component [99]. As a result, approximately 10% of the total number of channels were marked in all subjects. Raw data were preprocessed by applying a bandpass filter with cutoff frequencies of 0.01∼0.2 Hz. Motion artifacts were corrected by a correlation-based signal improvement algorithm [100].

### Behavior analysis

Trials with reaction times exceeding the threshold range (less than 0.4 s or greater than 1.5 s) are excluded from the analysis. The control and scope score of attention during vWM were calculated using the formula as follows:

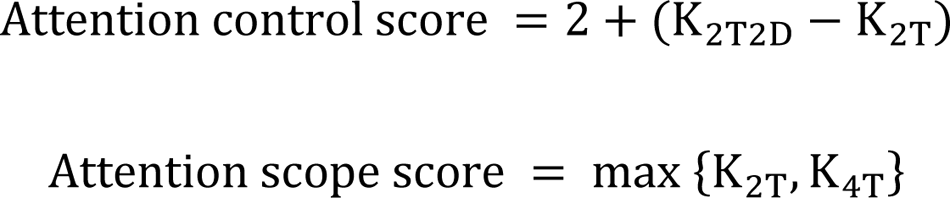

The dynamic K value was computed for the t th miniblock of the whole task, and its calculation formula is as follows:

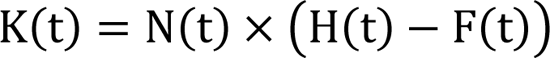

where N is the set size of the t th miniblock, t denotes the serial number of the miniblock, H(t) is the observed hit rate of the t th miniblock, and F(t) is the false alarm rate of the t th miniblock. The use of such formulas is based on the premise that during a certain period, an individual has a stabilized ability to hold in K items from an array of N items in vWM, which leads to correct performance on K/N of the trials.

### ERP analysis

We focused on the CDA component induced by the memory display. The baseline correction was calculated by subtracting the mean signal within the prior 200 ms to memory display onset. ERP waveforms were created by averaging all trials for six conditions (two visual hemifields × three experimental conditions). To isolate the CDA difference waves from the nonlateralized ERPs, the waveforms at the electrodes P3/4 and P7/8 ipsilateral to the side of the to-be-remembered items were subtracted from the contralateral ERPs. In this task, the memory display could induce a lateralized N2 posterior contralateral (N2pc) before the CDA component[101]. To measure CDA amplitude alone without N2pc contamination, the detection window stemmed from 500 ms, when the N2pc had ordinarily terminated.

During ERP analysis for two WM characteristics, the difference in CDA between baseline (2T) and the scope-increased conditions (4T) conditions was calculated as WM scope with t-statistical analysis. The differences between baseline (2T) and the control increased (2T2D) conditions were used to identify WM control processes. For each comparison, a test was calculated for each time point in the difference with random permutations (N = 1000). Our electrodes of interest were bilateral occipitoparietal electrodes (P3/4 and P7/8) based on previous findings [102].

### EEG oscillation

For the EEG oscillation analysis, artifact-free trials were transformed using a Morlet wavelet decomposition in 1 Hz steps as implemented by the MATLAB Fieldtrip package [103].

Subsequently, to calculate the induced neural oscillation responses without the evoked potential mask[104], the trial-average ERPs in the time domain were subtracted from the raw EEG signals on a trial-by-trial basis. Theta power changes (4–7 Hz) during retention were calculated to examine the scope-related oscillatory activity between baseline (2T) and scope-increased conditions (4T). The changes in control-dependent oscillatory activity were calculated to examine the difference between the baseline (2T) and the control-increased condition (2T2D). The theta band ERS measures were in line with the procedure reported [32, 33] as follows: 1) the reference interval (–400 to 0 ms) before memory display onset was subtracted from the power estimates at each time point, and 2) the difference was divided by the reference interval. Changes concerning baseline over time were referred to as ERS. Our electrodes of interest were middle frontal electrodes (FP1/2, F3/4, F1/2, and F_Z_).

### General linear model

We used a GLM to analyze hemodynamic response functions (HRFs) for different stimuli.

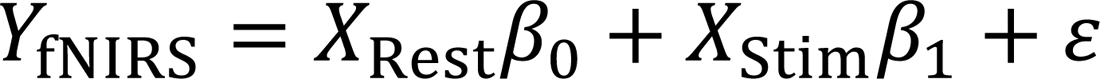

Where *Y*_fNIRS_ is the time series of the fNIRS signal, β_1_ is the regression coefficients, and ε is the error term. The design matrix *X*_stim_ is a convolution of the stimulus sequence, *X*_Rest_ is a convolution of the active rest state sequence, and basis functions and uses the same bandpass filter as was used for preprocessing. Basis functions are a set of Gaussian functions with widths of 2 s and steps of 2 s. Therefore, estimated HRF can be estimated using the following formula:

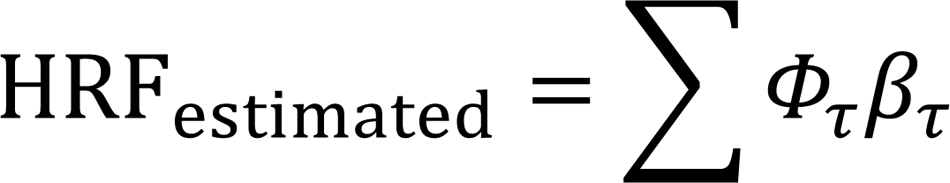

Where Φ_τ_ are the Gaussian basis functions and β_τ_ are regression coefficients associated with each basis function, with ττ representing the time delay (–2 s ∼ 8 s).

### Task-related activation

To evaluate the activation strength of hemodynamic activity in various conditions, a design matrix was constructed by convolving the cue events with a sequence of basis functions. These pivotal cue events prompt participants to actively engage in a cognitive task where they are required to memorize the visual content of stimuli occupying precisely half of the screen’s spatial area, as directed by arrow cues. The cue events mark critical moments in the experimental protocol, representing instances when participants are prompted to shift their attention and encode specific visual information. The basis functions are a series of Gaussian functions [93] with a width of 2 s and a step size of 2 s. A GLM analysis was then utilized to assess the hemodynamic activation levels.

T-test was conducted to identify the channels that rejected the null hypothesis β_4*T*_ = β_2*T*_ or β_2*T*_ = β_2*T*2*D*_ (p < 0.05) significantly:

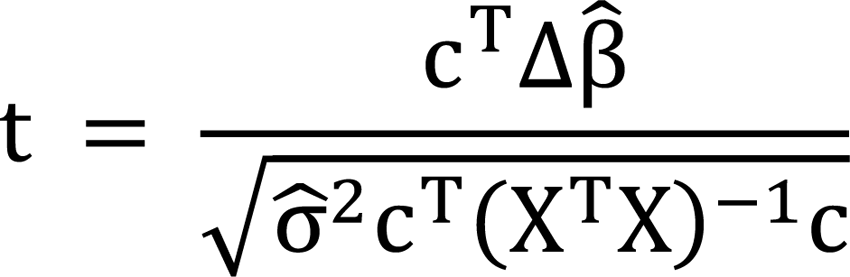

Where the estimate of weights Δβ- = β_4*T*_ − β_2*T*_ or β_2*T*2*D*_ − β_2*T*_, σ^2^ is the residual sum-of-squares divided by the degrees of freedom, c is the channel vector.

### Hemodynamic contralateral delay activity

To conduct an analysis similar to the typical approach in CDA involving lateralized analysis, we contrasted each channel of one hemisphere with its corresponding symmetrical channel in the opposite hemisphere. The initial step involves separately fitting and estimating the hemodynamic responses to stimulus events on both the same side and the opposite side. Subsequently, the relative changes in blood oxygen concentration (hemodynamic CDA) are obtained by subtracting the hemodynamic response on the ipsilateral side from the corresponding channel’s response on the contralateral side.

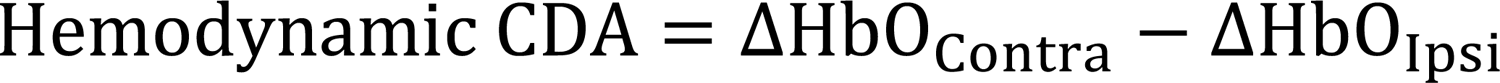

### EEG-informed analysis

In the present study, the design matrix contained three modeled hemodynamic response functions corresponding to *X*_Stim_, *X*_EEG_, and *X*_Rest_. In order to find NVC results that are specifically related to the EEG features and not to some general feature of the stimulus events, we used Schmidt-Gram orthogonalization. The orthogonalized sequences were then convolved with a standardized hemodynamic response function (HRF) convolved with a canonical hemodynamic response function, and subsequently modeled using the GLM to establish a linear relationship between EEG features and ΔHbO features.

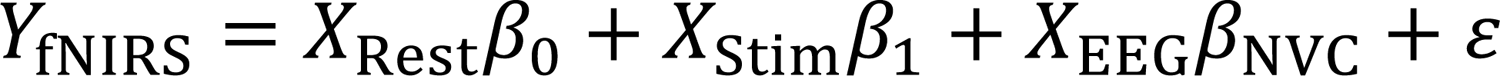

Where *y*_ΔHbO_ is the sequences of the ΔHbO features, *X*_EEG_ is the design matrix of EEG features, β_NVC_ is the regression coefficients to estimate NVC strengths, and ε is the error term.

*X*_Stim_ was orthogonalized with respect to *X*_EEG_ removing that part of *X*_Stim_ which was correlated to *X*_EEG_

### Modeling analysis

To assess the relationship between dynamic behavior and dynamic NVC in miniblocks at the group level, a linear mixed model was used with NVC as a predictor and K value as a response variable. Separate models were fit for each channel and estimates for group-level fixed effects (β_1_), intercepts (β_0_), and random effects (b_0i_, b_1i_) were obtained.

As the K value on 1st miniblock and the relationship between dynamic K and dynamic NVC may vary with each patient, a random intercept (b_0i_) and slope (b_1i_) were used.

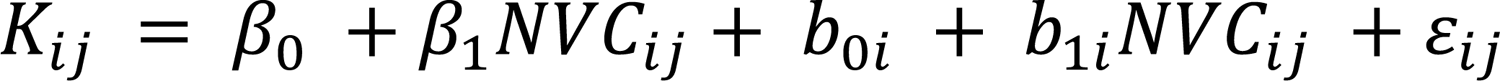

where b_0i_ and b_1i_ are random intercepts and slopes for subject i and *b*_0*i*_∼ *N*(0, τ^2^), *b*_1*i*_ ∼ *N*(0, τ_1*i*_ ^2^), K_ij_ denotes the jth fNIRS channel for subject i, NVC_ij_ indexes the EEG-informed NVC from j channel for subject i. The ANOVA analysis identified the relationship between dynamic K and dynamic NVC, and the significant main effects of NVC are reported after adjusting for false discovery rate (FDR) of 0.05. The t-statistic was used to determine the significance of ROI across the whole brain map.

### Statistical analysis

Bayes factor analyses with default priors (r = 0.707) were performed on the fNIRS and EEG data. Bayesian analysis was conducted using JASP software v. 0.9.2.0 to test the null hypothesis (BF_10_ < 0.333: substantial evidence for the null hypothesis; BF_10_ >1: support for H_1_ over H_0_).

In this study, ROC analyses were performed for the GLM binomial model. Each model was used to predict the attention performance based on EEG, ΔHbO, and NVC metrics, for each miniblock at a single subject level. We calculated the false positive rate (FPR, 1-specificity) and true positive rate (TPR, sensitivity) and generated the ROC by varying the p-value threshold (0 to 1). The AUC of ROC can be calculated to quantify the overall performance of each model. The higher AUC denotes that the model has a better performance for predicting behavior. To compare the performance of different models, a one-way repeated ANOVA test was employed to evaluate the difference in these metrics among the models based on EEG, ΔHbO, and NVC metrics.

## Notes

### Competing Interest Statement

The authors have declared no competing interest.

