## Supplementary Materials for "The neurovascular coupling in the attention during visual working memory"

**Zhang *et al.***

**Supplementary Information**

In traditional block-based experimental designs, the issue of overlapped blood oxygenation response signals often arises, leading to a reduction in the interpretative efficiency of the general linear model (GLM). This occurs due to the classical hemodynamic response function being based on an ideal model of short-term, single-event responses. The aim was to facilitate a rapid return of blood oxygenation signals to a consistently lower level, effectively mitigating mutual interference among miniblocks caused by continuous stimuli.

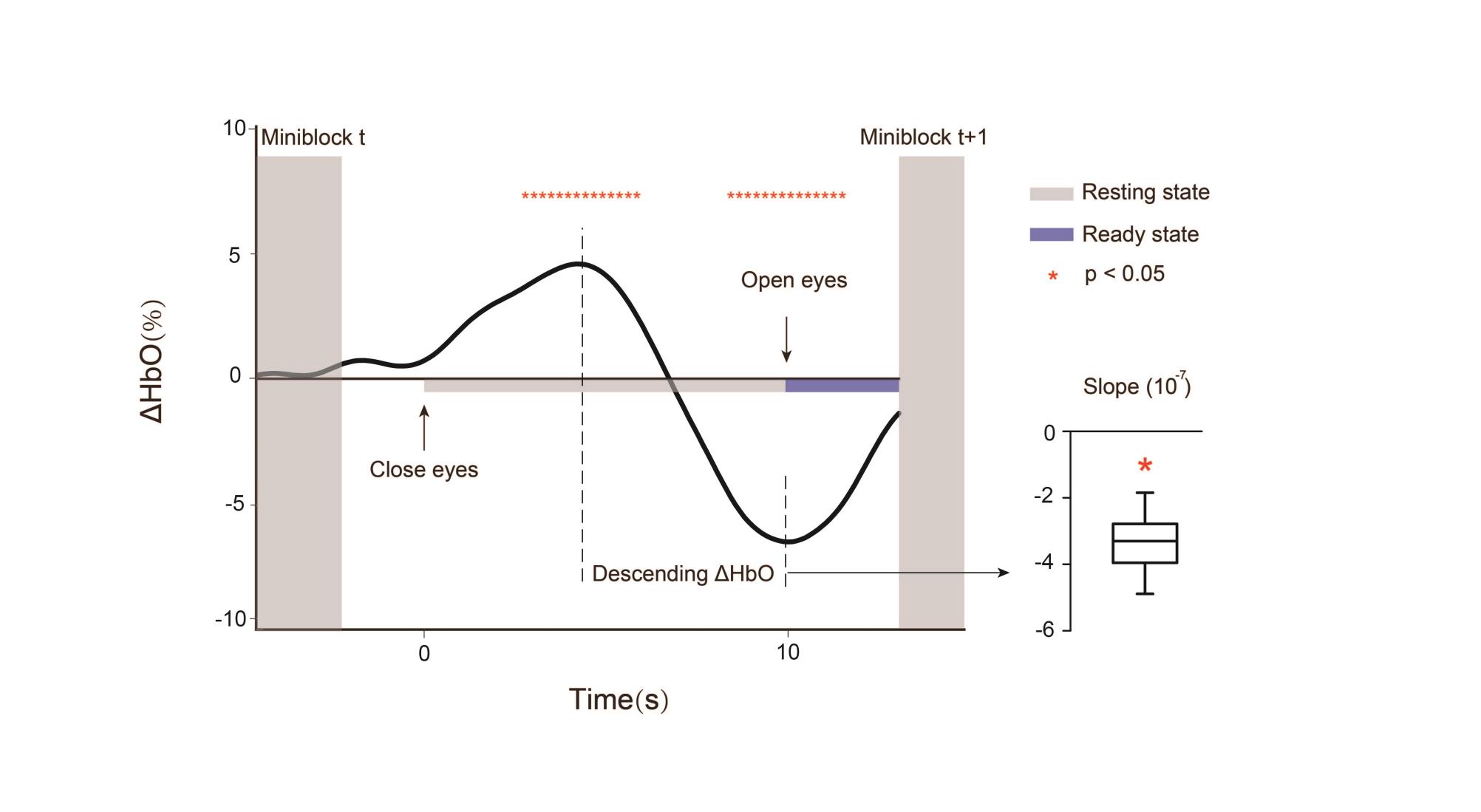

**Figure S1.** Temporal characteristics of ΔHbO signals during intertask resting periods. (A) During intervals between two miniblocks, a 10-second resting period was introduced, and participants were instructed to close or open their eyes at the boundary moments between the dark gray and light gray segments. The yellow segment indicates the countdown phase (approximately 2.2 s) before the formal initiation of the task. The dashed box highlights the interval from the peak of the ΔHbO response to the end of the resting phase. (B) Mean derivative values of the ΔHbO signal within this interval consistently exhibit a statistically significant negative trend.

As illustrated in Figure S1.A, the mean values of all channels are depicted for three different conditions, and the HbO variations during the resting phase exhibited a remarkably consistent trend. Following the conclusion of each miniblock, an immediate transition into a closed-eye resting state occurs. The ΔHbO signal exhibits a distinctive pattern: a gradual rise, reaching its peak at approximately 6 s, followed by a rapid decline until the onset of the eye-opening event. After reaching its peak following the peak of the ΔHbO signal and before the occurrence of the open-eye event, the statistical results within this timeframe demonstrated a significant decreasing trend in the ΔHbO signal, as illustrated in Figure S1.B. The above results indicated that introducing brief periods of rest indeed accelerates the signal decline and restores the baseline at a global level.

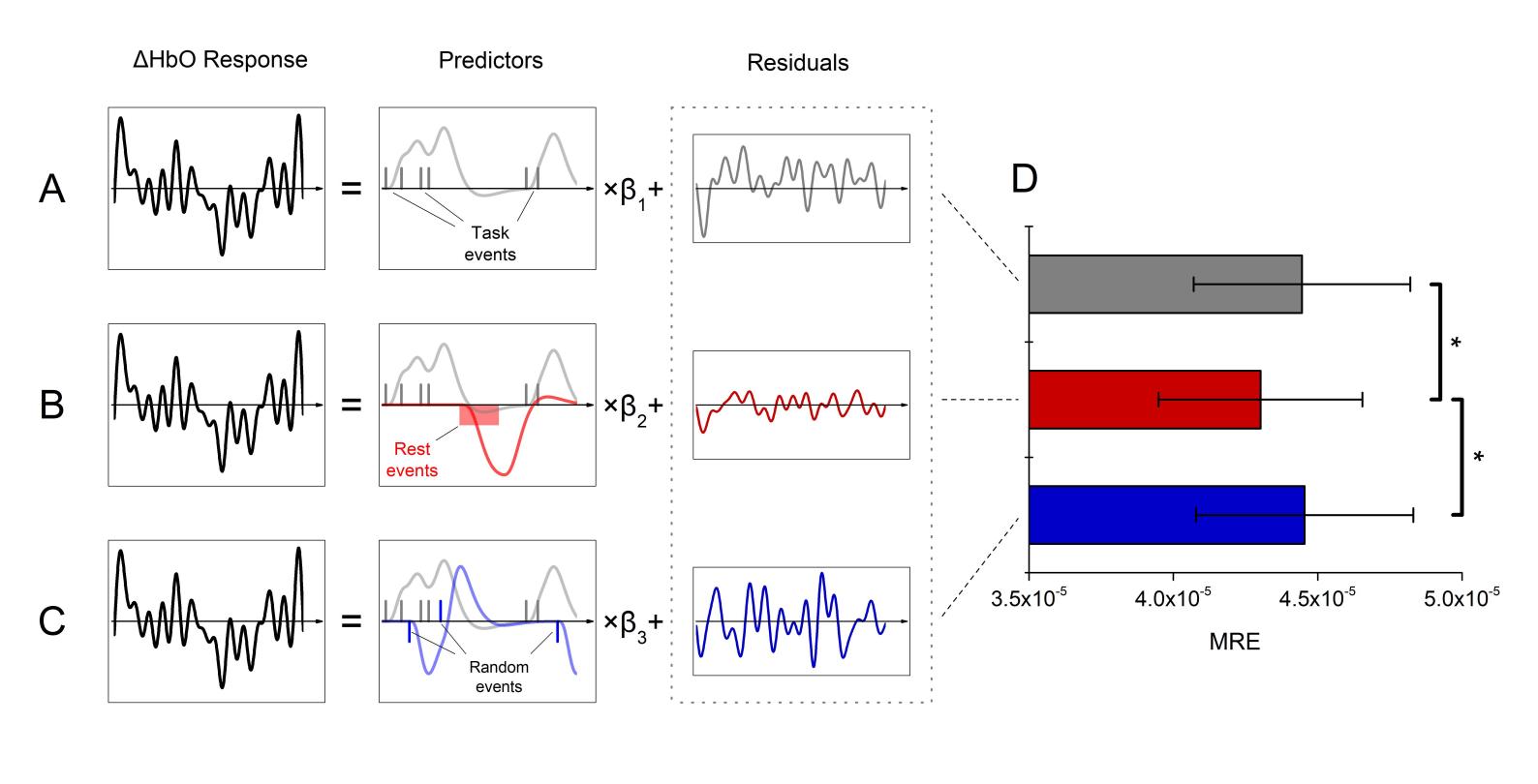

**Figure S2.** Impact of rest between miniblocks on ΔHbO responses in GLM analysis. (A) GLM fits with task-related events alone. (B) GLM fitting with task-related events and the influence of rest periods added. (C) GLM fitting with task-related events and the inclusion of random events to assess overfitting. (D) Compared to the other GLM configurations, the inclusion of rest periods significantly reduced the standard deviation (SD) of residuals associated with the ΔHbO response. However, the incorporation of random events had no significant effect on the SD of residuals.

To validate the effectiveness of the added resting period, not only in expediting signal recovery but also in enhancing the interpretability of the GLM, thereby bolstering the credibility of the results, we performed GLM estimations using distinct predictors (design matrices), as illustrated in Figure S2.A-C. We conducted three identical GLM estimations on all data sets, with the sole difference being the design matrix employed.

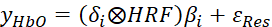

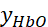
 is the signal for changes in blood oxygen concentration.
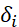
 are time series associated with different event conditions, where i=1 including only the task time; i=2 adding rest events: adding random event.
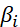
 corresponds to regression coefficients for different event combinations.
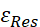
 is the residual term, and its standard deviation (SD) can reflect the explanatory power of the GLM, a lower SD indicating that the model fits the data better.

The first design matrix exclusively encompassed stimulus events; the second incorporated resting events; and the third introduced a series of random events. The metric was defined as the mean residual error (MRE) of the GLMs. MRE indicated the goodness of fitting between the expected model and the actual measured data. A smaller MRE would therefore represent a better fitting of the linear model. The residual statistics presented in Figure S2.D indicate that there is no significant difference in GLM residuals between task events and the addition of random events. However, the incorporation of resting events significantly reduces the residuals of the GLM fit compared to both scenarios. This to some extent suggested that adding resting periods indeed improves the model's explanatory capacity.

Furthermore, we investigated dispersion parameters and obtained corroborating results. Our findings from a model interpretative standpoint affirm our hypothesis that a miniblock-based experimental design effectively mitigates the interference stemming from overlapped consecutive stimulus events. This study contributes valuable insights toward refining the experimental design for enhanced accuracy in data interpretation.

**Table S1.**

| ROI | CH | MNI coordinates (X Y Z) | | | Anatomical label | Brodmann Area  (Talairach daemon) |
| --- | --- | --- | --- | --- | --- | --- |
| Control | 31 | 19 | 41 | 54 | Frontal_Sup_R (0.836) | Frontal eye fields (1.000) |
|  | 36 | 31 | 27 | 59 |  |  |
| Scope | 32 | 39 | 36 | 47 | Frontal_Mid_R (1.000) | Frontal eye fields (0.727) |
|  | 36 | 31 | 27 | 59 |  |  |

Numbers within brackets in the table denote the percentage of overlap

**Table S2.**

| ROI | CH | MNI coordinates (X Y Z) | | | Anatomical label | Brodmann Area  (Talairach daemon) |
| --- | --- | --- | --- | --- | --- | --- |
| Control | 10/20 | −25/25 | −84/−84 | 49/49 | Parietal_Sup (0.894) | Somatosensory Association Cortex, (0.618) |
| Scope | 10/20 | −25/25 | −84/−84 | 49/49 | Parietal_Sup (1.000) | Angular gyrus (0.968) |
|  | 9/21 | −51/51 | −75/−75 | 33/33 |  |  |

Numbers within brackets in the table denote the percentage of overlap
**Table S3.**

| ROI | CH | MNI coordinates (X Y Z) | | | Anatomical label | Brodmann Area  (Talairach daemon) |
| --- | --- | --- | --- | --- | --- | --- |
| Control  Theta | 37 | 49 | 20 | 50 | Frontal_Mid_R (0.987) | Frontal eye fields  (0.804) |
|  | 20 | 25 | −84 | 49 | Parietal_Sup_R (1.000) | Somatosensory Association Cortex (0.786) |
| Control  CDA | 6 | −61 | −60 | 33 | Parietal_Sup_L (0.671) | Angular gyrus  (0.509) |
|  | 31 | 19 | 41 | 53 | Frontal_Sup (0.702) | Frontal eye fields (0.870) |
|  | 35 | 7 | 31 | 64 |  |  |
|  | 40 | 19 | 12 | 71 |  |  |
|  | 37 | 49 | 20 | 50 | Frontal_Mid_R  （0.702） | Dorsolateral prefrontal cortex  (0.442) |
|  | 33 | 57 | 27 | 31 |  |  |
|  | 20 | −57 | −70 | 24 | Occipital_Sup_R  (0.439) | Somatosensory Association Cortex (0.646) |
|  | 21 | 51 | −75 | 33 |  |  |
|  | 19 | 61 | −60 | 33 |  |  |
| Scope  Theta | 39 | −7 | 15 | 72 | Frontal_Mid (0.732) | Frontal eye fields  (0.804) |
| Scope  CDA | 10 | −25 | −84 | 49 | Parietal_Sup_L (0.894) | Somatosensory Association Cortex, (0.618) |

Numbers within brackets in the table denote the percentage of overlap
